# SnRK2-mediated phosphorylation of SIZ1 enhances global SUMOylation under osmotic stress in Arabidopsis

**DOI:** 10.1101/2023.04.25.538284

**Authors:** Tian Sang, Bei Jia, Pengcheng Wang

## Abstract

SUMOylation is a highly dynamic posttranslational modification that plays a critical role in regulating plant stress responses. The global SUMOylation is quickly induced by dehydration and hyperosmotic stresses in plants, while the detailed mechanism underlying such SUMOylation dynamics is largely unknown. Here, we report that the SNF1-related protein kinase 2 (SnRK2) and SUMO E3 ligase SIZ1 module is crucial for the stress-induced increment of SUMOylation in Arabidopsis. Under osmotic stress, or application of phytohormone Abscisic Acid (ABA), the rapidly activated SnRK2s physically interact with and phosphorylate SIZ1, enhancing its stability. The Ser820 residue in C-terminal region of SIZ1 proteins is a functional SnRK2 phosphosite, whose phosphorylation is abolished in the high-order mutant of *SnRK2s*. The non-phosphorylatable SIZ1^S820A^ is unstable both *in vivo* and *in vitro*. We also noticed the degradation of SIZ1 is largely darkness-dependent, interestingly, independent of COP1, a key ubiquitin E3 ligase regulating photomorphogenesis. Multiple SUMOylation, Ubiquitination, and phosphorylation sites in SIZ1 proteins, which may coordinate the dynamics of SIZ1 proteins and global SUMOylation upon environmental changes. Our findings highlight the critical role of the SnRK2-SIZ1 module in regulating SUMOylation dynamics during plant stress responses and provide new insights into the regulatory mechanisms underlying this essential posttranslational modification.

## Introduction

Small ubiquitin-like modifier (SUMO) modification is a universal posttranslational modifications (PTMs) in eukaryotes. By covalently attaching the SUMO peptide to target proteins, SUMOylation controls the activity, localization, and turnover of many intracellular effectors. At least 20% of proteins in the human proteome could be modified by SUMOylation (Hendriks et al., 2017). In Arabidopsis, more than 1,000 proteins have been identified as putative SUMOylation targets in SUMOylome analyses (Rytz et al., 2018). The known SUMO-conjugated proteins are involved in transcription, epigenetic modification, hormone signaling, light response, nutrition, immunity, and environmental stress adaptations (Kong et al., 2020; Lee et al., 2007; Miura and Hasegawa, 2008; Miura et al., 2009; Park et al., 2011a; Richard et al., 2017; Sadanandom et al., 2015; Wotton et al., 2017). SUMOylation in Arabidopsis mainly depends on an enzyme cascade consisting of two SUMO activating enzymes (SAE), SAE1a and SAE1b, one SAE2, one SUMO conjugating enzyme 1(SCE1), and two E3 type SUMO ligases, SAP and MIZ1-1 (SIZ1) and METHYL METHANESULFONATE-SENSITIVE-21 (MMS21)/HIGH PLOIDY-2 (HPY2) (Park et al., 2011b). The mature SUMO is activated by a heterodimeric E1 consisting of SAE2 and one of two SAE1s. Then, the activated SUMO is transferred to SCE1 and conjugated to target proteins with the help of E3 ligases (Pichler et al., 2017; Streich and Lima, 2016). The SUMO conjugates can be removed by SUMO-specific proteases (Hickey et al., 2012). Mutants for the SUMO cascade enzymes *sae1*, *sae2,* and *sce1* single mutants, and *sumo1sumo2*, *siz1hpy2* double mutants are embryo lethal, suggesting an essential role of SUMOylation in plants (Ishida et al., 2012; Saracco et al., 2007).

Although SUMOylation can occur independently of SUMO E3 ligases *in vitro* (Bernier-Villamor et al., 2002), recent studies have revealed the essential role of SUMO E3 ligases in several physiological and developmental processes. The HPY2/MMS21 is involved in several developmental processes (Ishida et al., 2009; Liu et al., 2017; Liu et al., 2016; Zhang et al., 2017). For example, it is required for DNA repair and transcriptional gene silencing. Loss of function of the HPY2/MMS21 results in a premature transition from the mitotic cycle to the endocycle, leading to severe dwarfism with defective meristems (Ishida et al., 2009). The roots of *hpy2/mms21* have reduced responses to exogenous cytokinin and abnormal cell proliferation (Huang et al., 2009). The *siz1* mutant exhibits altered expression of brassinosteroid(BR) biosynthesis and signaling genes, resulting in smaller plant size than wild type plants. Additionally, SIZ1 is involved in regulating the FLC chromatin structure, which affects flowering (Jin et al., 2008). SIZ1 also plays a role in regulating nutritional responses to nitrogen and phosphate by facilitating nitrate reductase and PHR1 SUMOylation, respectively (Miura et al., 2005; Park et al., 2011a).

Besides regulating of growth and development, SUMO E3 ligases also play a crucial role in plant response to environmental changes. Various stresses like drought, heat stress, or reactive oxygen species (ROS) burst quickly increase SUMO conjugation, suggesting an enhanced SUMO E3 ligase activity upon these conditions (Li et al., 2017). SIZ1 has been known to function in heat stress, cold stress, drought, salt, and copper tolerance, as well as in plant immunity (Catala et al., 2007; Chen et al., 2011; Lee et al., 2007; Miura et al., 2007; Song Zhang and Meng, 2017; Yoo et al., 2006). Recent studies suggested that SIZ1 degradation is regulated by ubiquitin E3 ligase Constitutively Photomorphogenic 1 (COP1), the key regulator of photomorphogenesis (Kim et al., 2016; Kim et al., 2017). The drought and salinity-induced SUMO conjugates are enhanced in *cop1-4* mutant or *COP1* overexpressing lines, suggesting a potential role of COP1 in the stress-regulated SUMO conjugations (Kim et al., 2016). However, how stresses regulate the SUMO E3 ligase activity is largely unknown.

SNF1-related protein kinase 2s (SnRK2s) are key components in osmotic stress and phytohormone abscisic acid (ABA) signaling (Lozano-Juste et al., 2020). In Arabidopsis, there are ten members in the SnRK2 family, among which SnRK2.2, SnRK2.3, and SnRK2.6, are quickly activated by ABA, while all SnRK2s, except SnRK2.9, are quickly activated by osmotic treatments like high salinity or PEG (Boudsocq et al., 2004). Clade A of PP2C phosphatases bind and inhibit SnRK2s in the absence of ABA (Cutler et al., 2009; Fujii et al., 2009; Ma et al., 2009; Park et al., 2009). Upon stress condition, ABA binds to its receptor Pyrabactin Resistance 1 (PYR1)/PYR1-Like (PYL)/Regulatory Component of ABA Receptor (RCAR) proteins, which then inhibits the activity of PP2C. Upon releasing from PP2C-mediated inhibition, SnRK2s are phosphorylated by Raf-like protein kinases and, therefore, activated SnRK2s further phosphorylate downstream substrate to activate stress responses (Lin et al., 2020; Lozano-Juste et al., 2020; Soma et al., 2020; Takahashi et al., 2020; Lin et al., 2021). It has been observed that dysfunction of all the ten SnRK2s in *snrk2-decuple* (*snrk2.1/2/3/4/5/6/7/8/9/10*) or B4 subgroup RAFs in *OK^130^-null* resulted in hypersensitivity to osmotic stresses, indicating the importance of SnRK2s in stress responses (Fujii et al., 2011; Lin et al., 2020). While SnRK2s are crucial for osmotic stress and ABA signaling, their involvement in regulating global SUMO conjugations is yet to be investigated.

In this study, we found that the increase in global SUMO conjugations under osmotic stress mainly is mainly due to the accumulation of SIZ1 proteins. SnRK2s physically interact with and phosphorylate SIZ1, and thereby enhancing its stability. Ser820 in SIZ1 is a unique functional SnRK2 phosphosite *in vivo*. The stress-induced increase in Ser820 phosphorylation is abolished in *snrk2.2/3/6* triple (*snrk2-triple*) or *snrk2-decuple* mutants. Despite COP1’s dominant role in SIZ1 degradation in darkness, the degradation of SIZ1^S820A^ protein is not affected by dysfunction of COP1 in *cop1-4 SIZ1^S820A^/siz1-2* mutant. Therefore, osmotic stresses stabilize SIZ1 by SnRK2-mediated phosphorylation. This study presents a previously unreported phospho-regulation of SUMO E3 ligase SIZ1 and sheds new light on the interplay between different posttranslational modifications, which is critical for plant stress responses.

## Results

### SIZ1 is essential for ABA-and osmotic stress-induced SUMOylation

Using immunoblot with an anti-SUMO1 antibody, we measured the global SUMO conjugations upon ABA, NaCl, and mannitol treatments. Consistent with the previous findings (Catala et al., 2007; Kim et al., 2016; Wang et al., 2023; Zhang et al. 2013), ABA and osmotic stresses rapidly increased the global SUMO conjugations in Col-0 wild type (Figure 1A and 1B). Interestingly, the accumulation of SUMO conjugations induced by mannitol was strongly abolished in the *siz1-2* mutant compared to that in the Col-0 wild type (Figure 1C lane 1-4, 1D). Knockout of *HYP2* in the *hyp2-2* mutant only slightly impaired the mannitol-induced global SUMO conjugations (Figure 1C lane 5,6, 1D). These data suggested that osmotic stress-and ABA-induced SUMO conjugates are mediated by the enzymatic pathway, which primarily depends on SUMO E3 ligase SIZ1.

**Figure 1.**
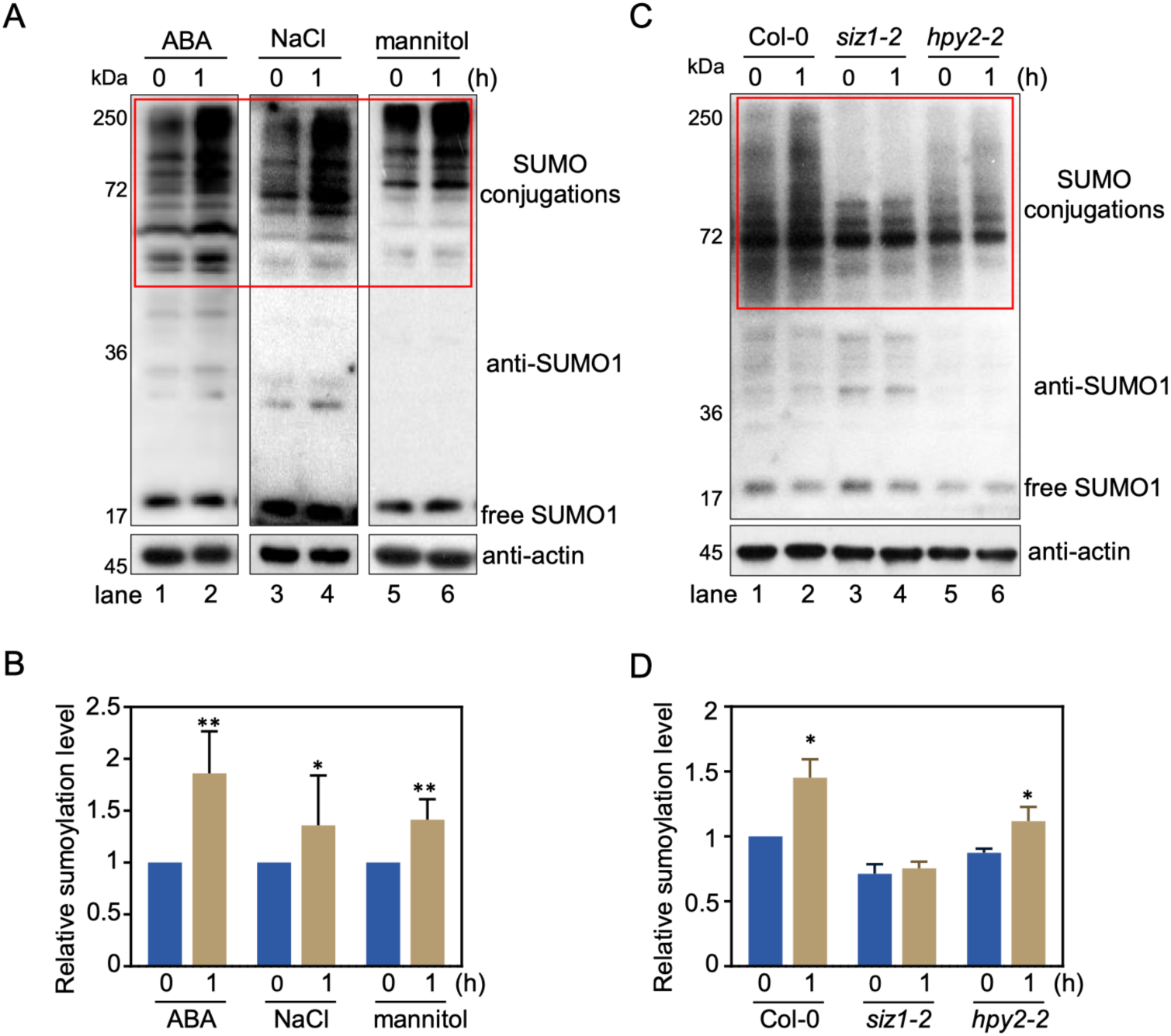
Osmotic stresses induce SUMO conjugations mainly depend on SIZ1. A. Accumulation of SUMO conjugations after indicated time of ABA (left panel), NaCl (middle panel), or mannitol (right panel) treatments. Immunoblot analysis with anti-SUMO1 (upper) or anti-actin (bottom) antibodies used to show the SUMO conjugates and loading of total proteins. The red box indicates the substrate band for SUMO binding. B. The relative SUMOylation levels from three independent assays in A (* p < 0.05, ** p < 0.01, Student’s t-test). C. Accumulation of SUMO conjugates after indicated time of mannitol treatment in *hpy2-2* and *siz1-2* mutants. Immunoblot analysis with anti-SUMO1 (upper) or anti-actin (bottom) antibodies used to show the SUMO conjugates and loading of total proteins. The red box indicates the substrate band for SUMO binding. D. The relative SUMOylation levels from three independent assays in C (* p < 0.05, Student’s t-test).

To evaluate the role of SIZ1-mediated SUMO conjugations in osmotic stress response, we assayed the gemination and seedling growth of *siz1-2* mutant on ABA and different osmotic stress treatment. As shown in Figures 2A and 2B, the seeds of *siz1-2* germinate slower than the wild type on either osmotic stress or ABA treatments. The *siz1-2* mutant also shows growth-arrested phenotypes upon various hyperosmotic stresses in the context of root growth (Figure 2C and 2D). These results suggested that SIZ1 is essential for osmotic stress-and ABA-induced SUMO conjugations and plays a crucial role in osmotic stress responses.

**Figure 2.**
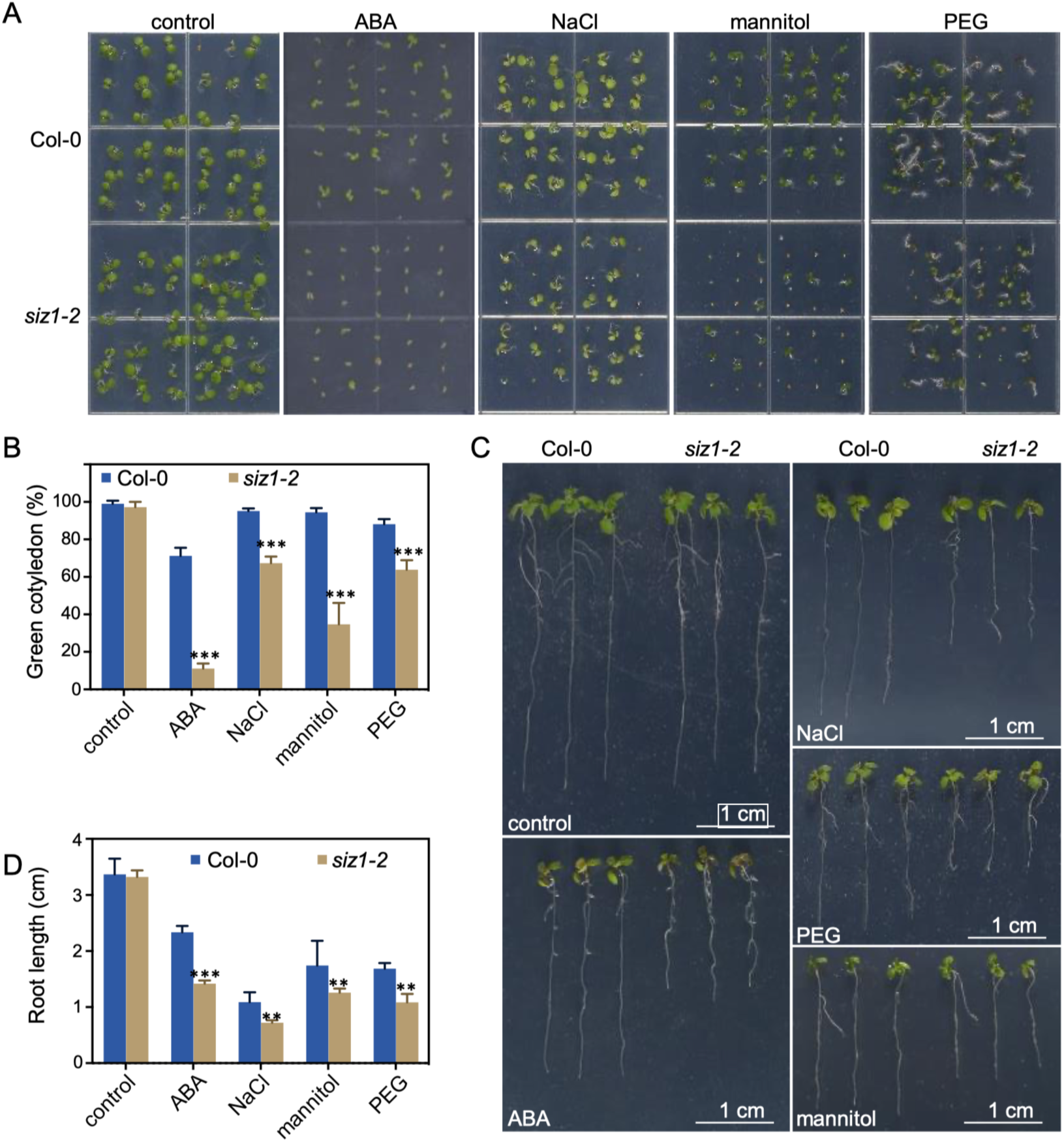
SIZ1 sensitive to ABA and osmotic stress. **A**. Photographs of seedlings after 5 days of germination on 1/2 MS medium containing indicated concentrations of 1 µM ABA, 150 mM NaCl, 250 mM mannitol and 0.4 mPa of PEG, respectively. **B**. The percentage of seedlings showing green cotyledons after 5 days of germination and growth on 1/2 MS medium containing ABA or different osmotic stresses. Values are mean± SD (n = 3; ***, *p* < 0.001. Student’s t-test). **C**. Photographs of seedlings growing 10 days after transfer to medium containing 15 µM ABA, 150 mM NaCl, 250 mM mannitol and 0.4 mPa of PEG, respectively. **D**. Quantitative measurement of the root length of the seedlings in (C). Values are mean ± SD (n = 15, **, *p* < 0.01; ***, *p* < 0.001. Student’s t-test).

### SnRK2s interact with and phosphorylate AtSIZ1

To study how ABA and osmotic stresses regulate SIZ1 function, we first measured the *SIZ1* mRNA levels under different osmotic stress and ABA treatments. Quantitative RT-PCR results showed that the expression of *SIZ1* was not induced by exogenous ABA and osmotic stress (Supplemental Figure 1A). By using an antibody specific to Arabidopsis SIZ1 protein (Supplemental Figure 1B), we found that the abundance of SIZ1 protein significantly increased under ABA or osmotic stress conditions (Figures 3A and 3B), consistent with the observed increase in SUMO-conjugation levels (Figure 1A). These data suggest that the accumulation of SIZ1 protein induced by osmotic stress and ABA may depend on post-translational regulation. Pre-incubation with MG132, an inhibitor of the proteasome-dependent protein degradation pathway, increased the SIZ1 levels over time (Figure 3C), indicating that ubiquitination and the 26S proteasome complex may contribute to the stability of SIZ1 proteins.

**Figure 3.**
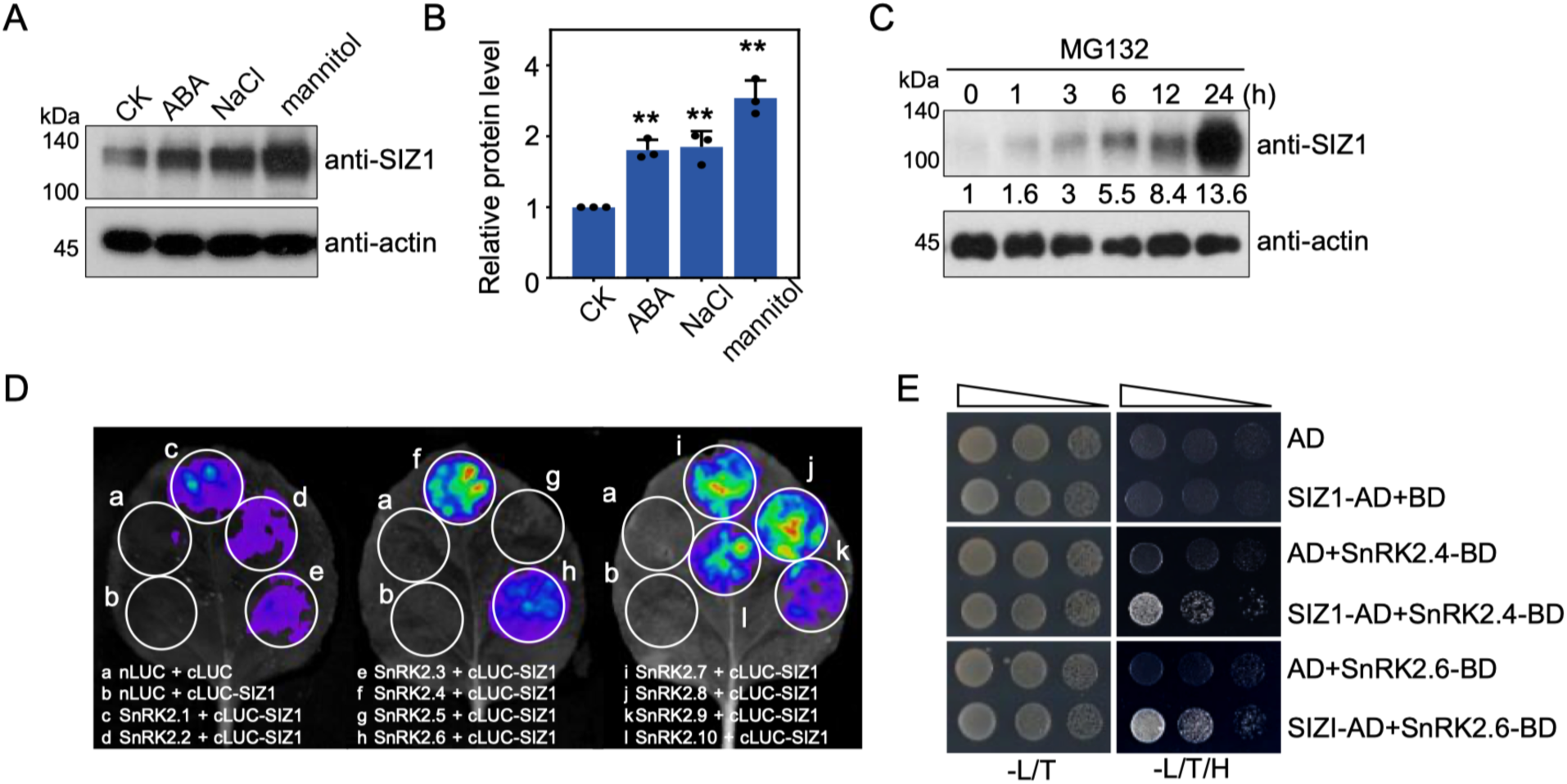
SnRK2s interact with SIZ1. **A**. The abundance of SIZ1 protein after indicated time of ABA, NaCl, or mannitol treatments. Immunoblot analysis with anti-actin antibody used to show loading of total proteins. **B.** The relative protein levels from three independent assay in A (**, p < 0.01. Student’s t-test). **C**. The abundance of SIZ1 protein after indicated time of 20 µM MG132 treatment in 10-day-old seedlings. Immunoblot analysis with anti-actin antibody used to show loading of total proteins. The numbers below anti-SIZ1 blots indicate the relative band intensities of SIZ1. The ratio of the first clear band was set to 1 for each blot. **D.** SIZ1 interacts with SnRK2s in split LUC complementation assay in *Nicotiana benthamiana* leaves. The nLUC/cLUC and nLUC/SIZ1-cLUC combination were used as negative controls. **E.** SnRK2.6 and SnRK2.4 interact with SIZ1 in a yeast-two-hybrid assay.

The ABA-and osmotic stress-activated SnRK2s are central components in stress responses (Boudsocq et al., 2004; Fujii et al., 2007, 2011). Therefore, we hypothesized that SnRK2s may also play a role in the stabilization of SIZ1 upon osmotic stresses. To validate this hypothesis, we investigated the interaction between SIZ1 and SnRK2s by the split-luciferase complementation (split-LUC) assay in tobacco leaves. Our result showed SIZ1 strongly interacts with SnRK2.4, SnRK2.7, SnRK2.8, and SnRK2.10, respectively, and shows weak interaction with SnRK2.1, SnRK2.2, SnRK2.3, SnRK2.6, and SnRK2.9 (Figure 3D). However, we did not detect the interaction between SnRK2.5 and SIZ1 in the split-LUC system, nor did we detect the interaction between n-LUC empty vector and SIZ1-cLUC, c-LUC empty vector and SnRK2s-nLUC, or n-LUC and c-LUC combinations (Supplemental Figure 2A). Yeast-two-hybrid assays also supported the physical interaction between SnRK2.4/SnRK2.6 and SIZ1 (Figure 3E).

Furthermore, we performed a bimolecular fluorescence complementation (BiFC) assay and found that YFP fluorescence signals were only detected in the nucleus of N. benthamiana cells that co-expressed SnRK2.4/6-YNE and SIZ1-YCE (Supplemental Figure 2B). Consistently, SIZ1-RFP was co-localized with SnRK2.4/6-GFP in nucleus of Arabidopsis protoplasts (Supplemental Figure 2C). The GST-pull down assay result showed that both GST-SnRK2.4 and GST-SnRK2.6, but not GST, could pull down MBP-SIZ1-C protein (Supplemental Figure 2D). Taken together, our results indicate that SnRK2s interact with the SIZ1 protein, supporting the hypothesis that SnRK2s may be involved in the stabilization of SIZ1 upon osmotic stresses.

### SnRK2s phosphorylate SIZ1 protein

Encouraged by the direct interaction between SnRK2s and SIZ1, we then determine whether SIZ1 is a substrate of SnRK2s. Due to fail to obtain enough amount of recombinant full-length SIZ1 protein, we used three SIZ1 fragments, SIZ1-N (aa1-150), SIZ1-M (aa59-537), and SIZ1-C (aa435-873) for *in vitro* kinase assay (Figure 4A). The results showed that recombinant SnRK2.4 strongly phosphorylates the SIZ1-C fragment, but only slightly phosphorylates SIZ1-M *in vitro* (Figures 4B and 4C). SnRK2.4 did not phosphorylate the N-terminal fragment SIZ1-N (Figure 4C). SnRK2.6 also can phosphorylate, though weaker than SnRK2.4, the SIZ1-C fragment (Supplemental Figure 3A), which is consistent with the weaker interaction between SnRK2.6 and SIZ1-C, than that between SnRK.4 and SIZ1-C (Supplemental Figure 2D). LC-MS/MS assay identified three putative phosphosites Ser794, Ser796, and Ser820, in SIZ1 after incubation with ^18^O-labeled ATP and SnRK2.4 (Figure 4D, Supplemental Figure 3B). Another phosphosite, Ser95 in SIZ1-M fragment, also present in the result of LC/MS-MS (Supplemental Figure 3C). Thus, SnRK2s phosphorylate multiple sites in SIZ1 protein.

**Figure 4.**
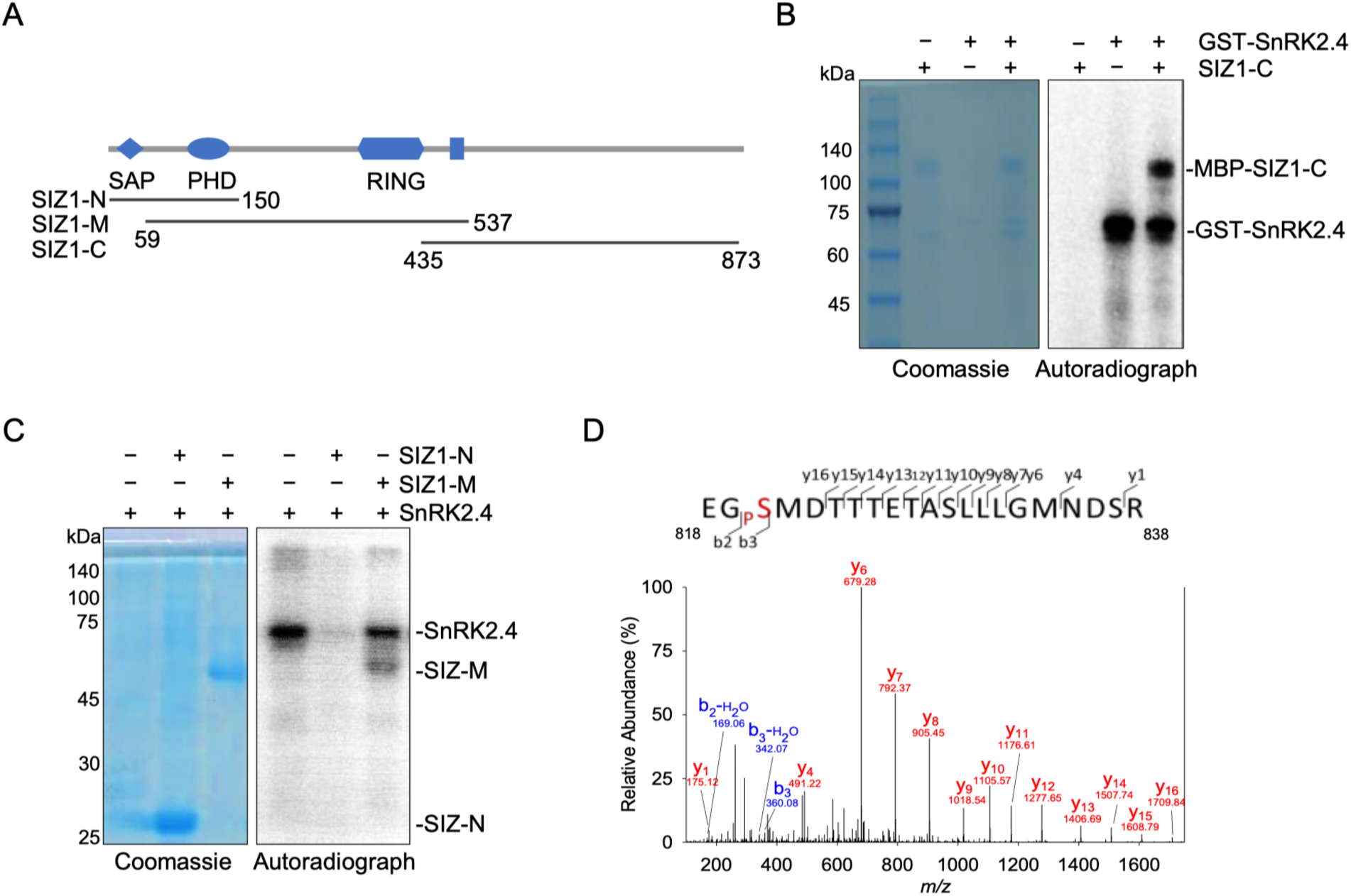
SIZ1^S820A^ is unstable *in vivo* and *in vitro*. **A**. Different SIZ1 fragments used in the in vitro kinase assay. **B.** Recombinant SnRK2.4 was used to phosphorylate recombinant SIZ1-C in the presence of [γ-^32^p] ATP. Autoradiograph (right) and Coomassie staining (left) show phosphorylation, loading of purified SIZ1-C and SnRK2.4. **C**. SnRK2.4 phosphorylates SIZ1-M in vitro. Recombinant SnRK2.4 was used to phosphorylate recombinant SIZ1-N and SIZ1-M in the presence of [γ-^32^p]ATP. Autoradiograph (left) and Coomassie staining (right) show phosphorylation, loading of purified SIZ1-N (aa 1-150), SIZ1-M (aa 59-537), and SnRK2.4. **D.** The MS/MS spectrum showing that the ^18^O-phosphopeptide EGpSMDTTTETASLLLGMNDSR contains the phosphoserines Ser820 in SIZ1.

Interesting, we also noticed that SnRK2.4 could phosphorylate the recombinant GST-HYP2 *in vitro* (Supplemental Figure 3D). As SIZ1 has a dominant role in stress-induced global SUMO conjugation, we only focused on the phosphor-regulation of SIZ1 in this study.

### Ser820 phosphorylation is required for AtSIZ1 stabilization

To verify these putative SnRK2s phosphosites *in vivo*, we introduced SIZ1 carrying the wild type, phosphomimic Ser (S) to Asp (D) mutation, and non-phosphorylatable Ser (S) to Ala (A) mutations of these four putative phosphosites under the control of the native promoter into the *siz1-2* mutant. Introducing the SIZ1^S95A^, SIZ1^S95D^, SIZ1^S794A^, SIZ1^S794D^, SIZ1^S796A^, SIZ1^S796D^, and SIZ1^S820D^ mutants, as well as the wild type SIZ1, successfully resorted the *siz1-2* phenotype regarding seedling size (Figure 5A-D and Supplemental Figure 4A-C). However, introducing SIZ1^S820A^ was unable to complement the phenotype of the *siz1-2* in the context of seeding growth or ABA sensitivity (Figure 5A and 5D). The *siz1-2/SIZ1^S820A^* seedlings displayed growth-arrested phenotype in the soil and ABA-sensitivity on the medium, which was comparable to the *siz1-2* mutant (Figure 5A, 5D and Supplement Figure 4C, 4F).

**Figure 5.**
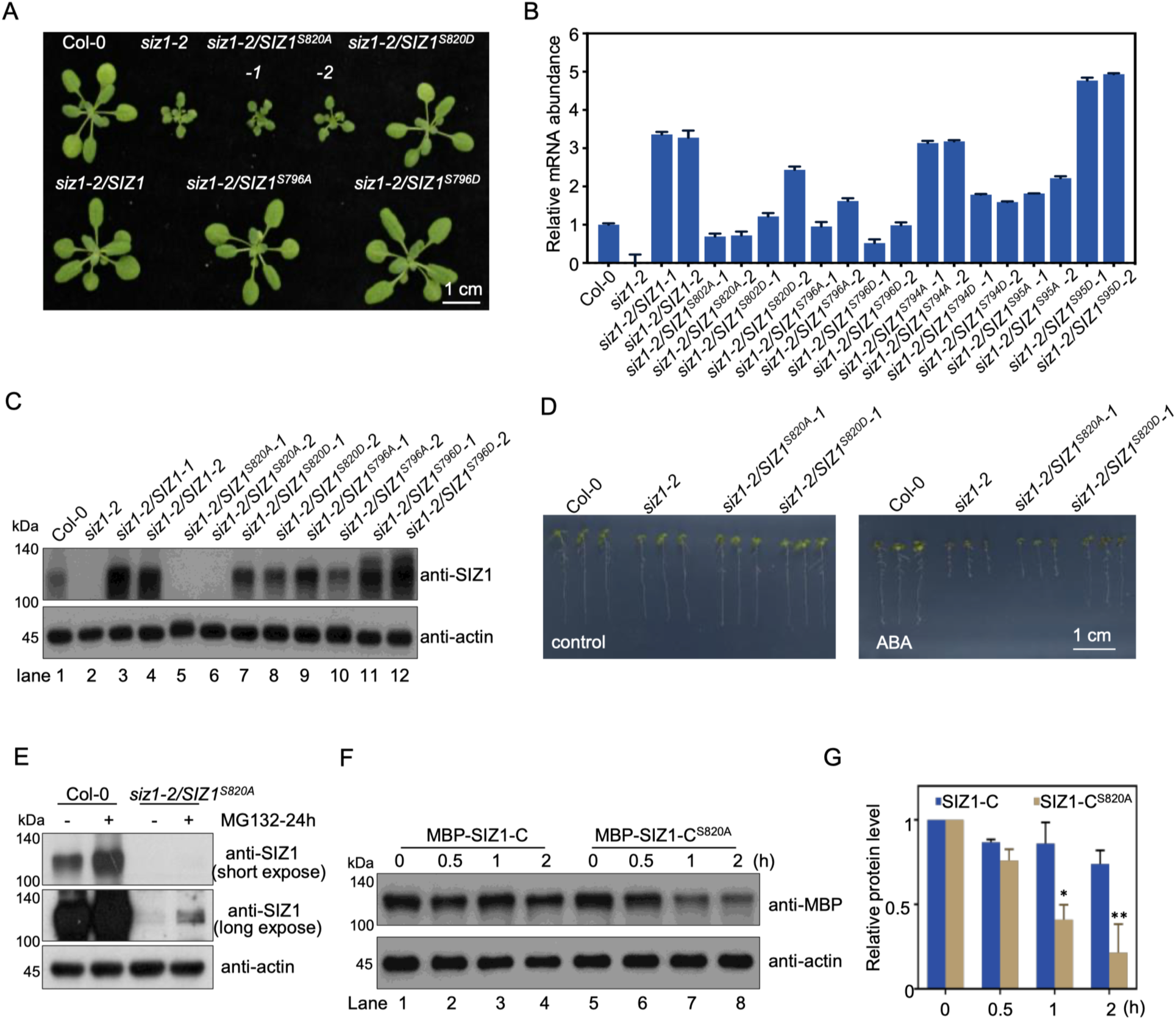
SIZ1^S820A^ is unstable *in vivo* and *in vitro*. **A**. Photographs of 4-week-old seedlings from the wild type, *siz1-2*, and transgenic plants. **B**. Transcription of SIZ1 genes in wild type, *siz1-2*, and transgenic plants. **C**. Immunoblot shows the SIZ1 abundance in wild type, *siz1-2*, and transgenic plants. Immunoblot with anti-actin antibody (bottom) used to show loading of total proteins in each lane. **D**. Photographs of seedlings growing 8 days after transfer to medium containing 20 µM ABA. **E**. Immunoblot shows the SIZ1 abundance in wild type and *siz1-2/SIZ1S820A* plants with or without MG132 treatment in 3-week-old plants leaf. Immunoblot with anti-SIZ1 antibody (upper and middle) used to show the SIZ1 abundance in short and long expose, respectively. Immunoblot with anti-actin antibody (bottom) used to show loading of total proteins in each lane. **F**. Immunoblot shows the MBP-SIZ1-C or MBP-SIZ1-CS820A abundance after incubation with the protein extract from 10-d-old wild type seedlings at indicated time. Immunoblot with anti-actin antibody (bottom) used to show loading of total proteins in each lane. **G**. Quantification analysis of (E). The band intensities were measured using the ImageJ tool. For each protein, the relative levels at 0 h were defined as 1. Error bars indicate SD from three independent experiments (* p < 0.05, ** p < 0.01, Student’s t-test).

Interestingly, despite comparable expression levels of the *SIZ1^S820A^*gene in *siz1-2/SIZ1^S820A^* lines compared to other *siz1-1/SIZ1* transgenic plants (Figure 5B and Supplement Figure 4D), there is no detectable SIZ1^S820A^ protein in the seven independent alleles of *siz1-2/SIZ1^S820A^* (Figure 5C, Supplemental Figure 4E). Furthermore, even after incubating with MG132 for 24 h, only a minimal amount of SIZ1^S820A^ proteins was detected in *SIZ1^S820A^/siz1-2* plants (Figure 5E). The stability of wild-type SIZ1 and SIZ1^S820A^ was also evaluated using the cell-free degradation assay, which showed that SIZ1^S820A^ is less stable than wild-type SIZ1 (Figure 5F and 5G). These results suggest that phosphorylation at Ser820 in the SIZ1 protein is essential for SIZ1 stabilization.

### ABA-and osmotic stress-induced Ser820 phosphorylation depends on SnRK2s

The mutation on the Ser820 residue strongly abolished SnRK2 phosphorylation on SIZ1-C in the in vitro kinase assay (Figure 6A), providing further support for the importance of this phosphosite. To further confirm the in vivo phosphorylation of Ser820 by SnRK2s, a parallel reaction monitoring (PRM) assay was used to measure the Ser820 phosphorylation in the wild-type and *snrk2-decuple* mutant seedlings, with or without mannitol or ABA treatment. The results showed that Ser820 phosphorylation is significantly induced by mannitol treatment in the wild-type seedlings (1.6-fold, p < 0.01, student’s *t*-test), while no change in the *snrk2-decuple* mutant seedlings (Figure 6B, upper panel). Similarly, ABA treatment also increases Ser820 phosphorylation (2.5-fold, p < 0.01, student’s *t*-test) in wild-type but not in *snrk2-triple* mutant seedlings (Figure 6C, bottom panel). Supplemental Figure 5A displays the extracted chromatograms of fragment ions of EGpSMDTTTETASLLLGMNDSR peptides, which contain the Ser820 phosphorylation. These findings suggest that the osmotic stress-and ABA-induced Ser820 phosphorylation is dependent on the presence of SnRK2s. To further investigate this, we measured the abundance of SIZ1 protein in WT and *SnRK2* high-order mutants by immune blot. As a high concentration ABA is used to isolate *snrk2-decuple* homozygous seedlings, the *snrk2-decuple* are unsuitable for further mannitol treatment. Nevertheless, we found that the dysfunction of ABA-activated SnRK2.2/2.3/2.6 completely abolished the ABA-induced accumulation of SIZ1 proteins in *snrk2-triple* mutant (Figure 6D, upper panel, lanes 6-8 compared to lane 5, Figure 6F). Interestingly, the basal level of SIZ1 protein abundance in *snrk2-triple* is slightly higher than that in Col-0 wild type (Figure 6D, upper panel, lanes 5-8 compared to lane 1), possibly due to an unknown feedback mechanism increasing SIZ1 expression in *snrk2-triple* (Supplemental Figure 5B). We also repeated the same assay using the wild-type and *snrk2-triple* seedlings germinated and grown in the dark, a condition that impairs SIZ1 stability (Kim et al., 2016). The results clearly showed that ABA induced the accumulation of SIZ1 proteins in wild-type seedlings, and the SIZ1 protein was undetectable in the *snrk2-triple* mutant (Figure 6E and 6G). Taken together, these data demonstrate that the Ser820 residue in SIZ1 is a functional SnRK2 phosphosite in plants, and its phosphorylation corresponds to the ABA-and osmotic stress-induced accumulation of SIZ1 proteins.

**Figure 6.**
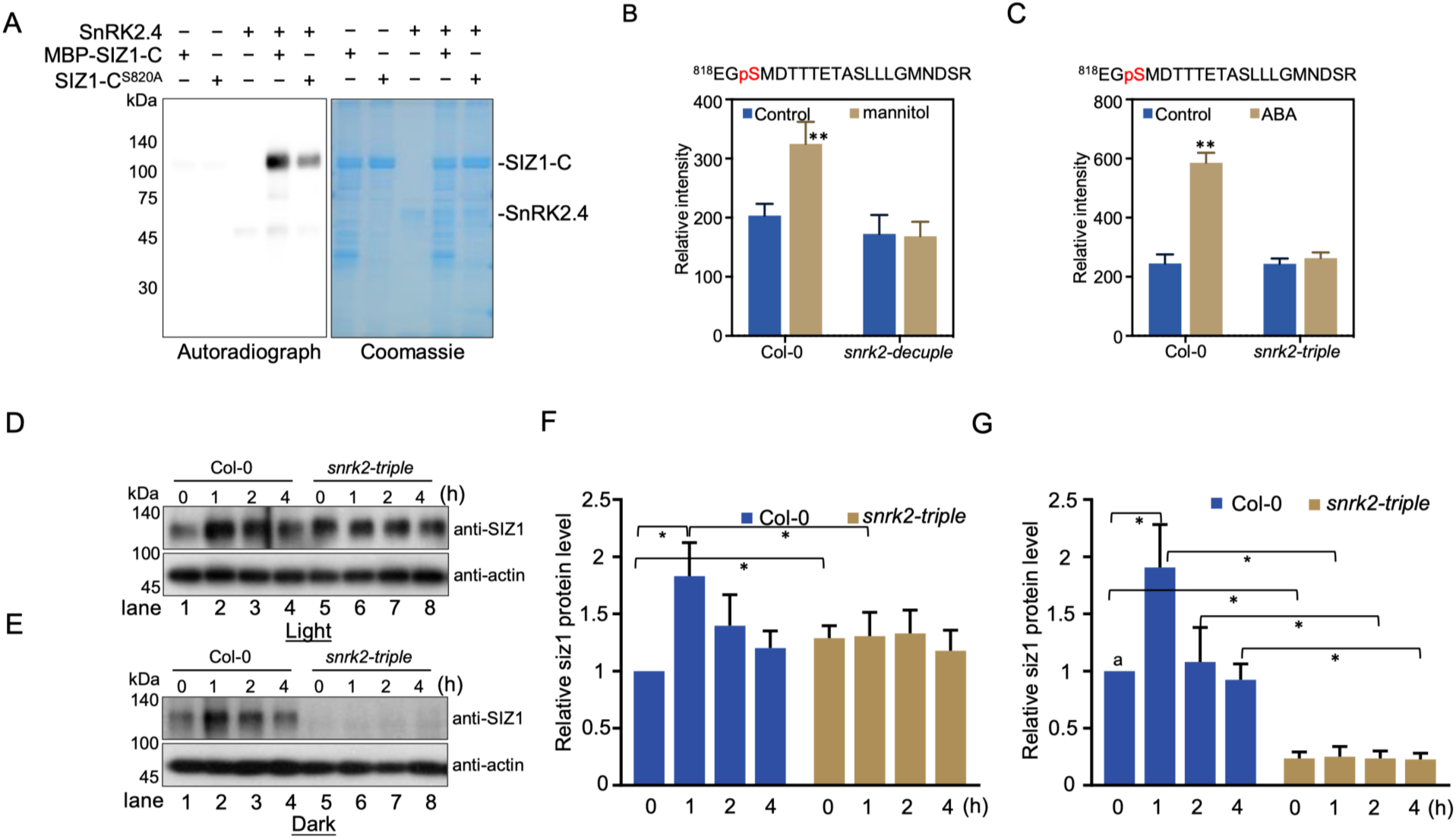
SnRK2 and dark coordinate SIZ1 stability. **A**. Ser820Ala mutation impaired SIZ1-C phosphorylation. Recombinant SnRK2.4 was used to phosphorylate recombinant SIZ1-C and SIZ1-CS820A in the presence of [γ-32p]ATP. Autoradiograph (left) and Coomassie staining (right) show phosphorylation, loading of purified SIZ1-C and SnRK2.4. **B** and **C**. The relative intensity of EGpSMDTTTETASLLLGMNDSR phosphopeptide in wild-type, *snrk2-triple*, and *snrk2-decuple* seedlings with or without ABA or mannitol treatment for 1h. ** p < 0.01 (Student’s t-test) for the indicated pairs of seedlings. **D.** SIZ1 abundance after indicate time of ABA treatment in Col-0 and *snrk2-triple* mutant under light condition. Immunoblot analysis with anti-actin antibodies used to show loading of total proteins in each lane. **E.** SIZ1 abundance after indicate time of ABA treatment in Col-0 and *snrk2-triple* mutant under dark condition. Immunoblot analysis with anti-actin antibodies used to show loading of total proteins in each lane. **F.** The relative protein levels from three independent assay in E (*, p < 0.05. Student’s t-test). **G.** The relative protein levels from three independent assay in F (*, p < 0.05. Student’s t-test).

### SIZ1^S820A^ degradation is independent on COP1

The differential performance of SIZ1 protein in dark and light conditions suggested a potential crosstalk between ABA/osmotic stress signaling and light responses. The key regulator of photomorphogenesis, COP1, has been known to regulate SIZ1 protein stability by ubiquitination (Kim et al., 2016). By crossing *cop1-4* and *siz1-2/SIZ1^S820A^-1*, we generated *cop1-4/siz1-2/SIZ1^S820A^*, a *cop1-4/siz1-2* double mutant carrying Ser820Ala mutated SIZ1. Interestingly, the SIZ1^S820A^ protein was still undetectable in *cop1-4/siz1-2/SIZ1^S820A^* (Figure 7A, lane 4 and 5). To further dissect the role of 26S proteosome pathway in SIZ1^S820A^ degradation, we also measure the SIZ1^S820A^ abundance after 12 h incubating of MG132. The result showed that inhibiting of 26S proteosome pathway by MG132, which increase wild type SIZ1 proteins in Col-0 wild type seedlings, didn’t affect the stability of SIZ1^S820A^ in either Col-0 or *cop1-4* (Figure 7A, lane 3 and 6). In addition, both wild type and Ser820Asp mutated SIZ proteins undergo degradation in darkness (Figure 7B). Although darkness and ABA/osmotic stress coordinately regulate the protein stability and abundance of SIZ1, the SIZ1^S820A^ degradation is likely independent on COP1, or 26S proteosome pathway.

**Figure 7.**
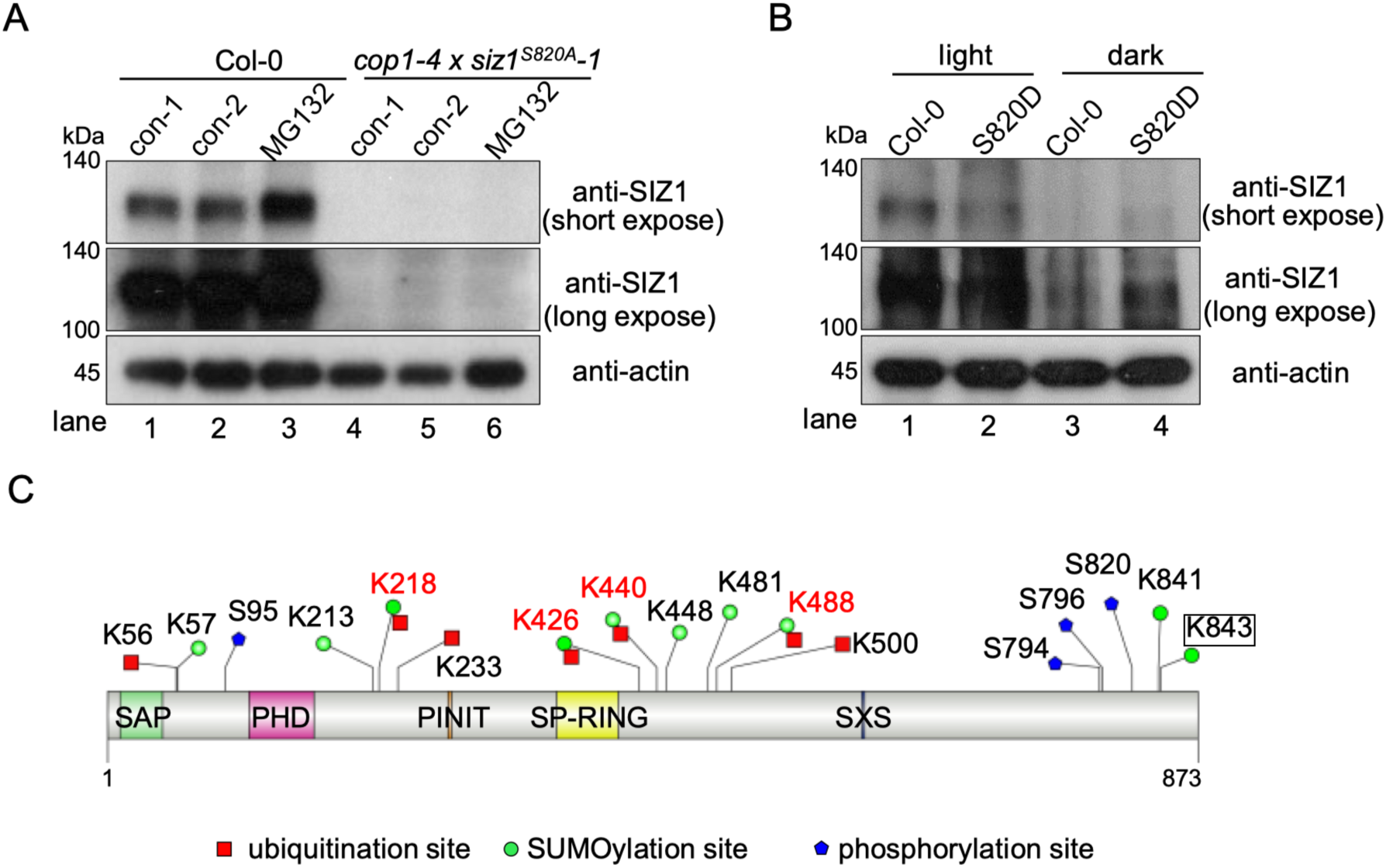
SIZ1^S820A^ protein stability is independent with COP1. **A**. SIZ1 abundance in *cop1-4 SIZ1^S820A^/siz1-2* with or without 12h MG132 treatment. **B.** SIZ1 abundance in wild type and *SIZ1^S820D^/siz1-2* plants under light and dark conditions. Immunoblot with anti-actin antibody used to show loading of total proteins in each lane. **C.** SUMOylation, ubiquitination, and phosphorylation sites in SIZ1 protein identified by mass spectrometry in His-H89R-SUM1 plants. Red square: Ubiquitination sites, green circle: SUMOylation sites, blue pentagon: Phosphorylation sites.

To further measure the PTM in SIZ1 protein upon ABA treatment, we identified the ubiquitination and SUMOylation sites in immunoprecipitated SIZ1 by liquid chromatography with tandem mass spectrometry (LC-MS/MS), using a *His-H89R-SUM1* transgenic plant (Miller et al., 2010) (Supplemental Figure 6A-H). There are seven ubiquitination sites (GG sites) and nine SUMOylation sites (QTGG sites) identified in immunoprecipitated SIZ1 proteins upon ABA treatment. Interestingly, four lysine residues, Lys218, Lys426, Lys440, and Lys488 could be modified by both SUMOylation and ubiquitination (Figure 7C, and Supplemental Dataset 1). Together with previous studies (Kim et al., 2106; Lin et al., 2006), our results suggested phosphorylation, ubiquitination and SUMOylation coordinately modulate stability of SIZ protein, to respond to dark/light and environmental changes.

## Discussion

It is well-documented that global SUMOylation increases in response to various environmental stresses. In Arabidopsis, one of two SUMO E3 ligases, SIZ1, plays a dominant role in multiple stress responses (Catala et al., 2007; Miura and Hasegawa, 2008; Miura et al., 2007). SIZ1 is known to regulate key transcription factors, such as ABI5, ICE1, MYB30, CaDSIZ1, and NF-YC10, to modulate the transcription of stress-responsive genes (Miura et al., 2007; Miura et al., 2009; Zheng et al., 2012; Joo et al., 2022; CHuang et al., 2023). Recently studies also suggest that SIZ1 also participates in chromatin remodeling and epigenetic regulation to modulate transcription in response to osmotic stresses or application of ABA (Miura et al., 2020; Han et al., 2020; Kong et al., 2020; Rytz et al., 2018). Despite its role in stress-induced SUMO conjugation, the precise mechanism by which SIZ1’s biological function is promoted by stress remains unclear. In this study, we report that SnRK2s, which are master regulators in osmotic stress and ABA signaling, phosphorylate and stabilize SIZ1 to promote global SUMOylation under stress conditions. SnRK2s mediate phosphorylation of SIZ1 at Ser820, which is essential for its stabilization. Abolishing SnRK2-mediated SIZ1 phosphorylation by Ser820Ala mutation or disrupting snrk2s in *snrk2-triple* plants under dark conditions led to undetectable levels of SIZ protein. Our study thus presents a mechanistic view of how SnRK2s kinase phosphorylates SIZ1 to increase its protein stability under osmotic stress conditions in Arabidopsis. While the activities of two SUMO E3 ligases, polycomb group protein Pc2 (Pc2), and Protein inhibitor of activated STAT1 (PIAS1), are known to be regulated by phosphorylation in animal cells (Liu et al., 2007; Swaminathan et al., 2004), phospho-regulation of SUMO E3 ligases by SnRK2-mediated phosphorylation has not been reported in plants before. Therefore, it appears that both animals and plants utilize similar phosphorylation regulation mechanisms to promote SUMO E3 ligase activity and global SUMOylation, and our study sheds new light on crosstalk between posttranslational modification (PTM) upon environmental stimuli in plants. Other environmental challenges, like high temperature, also dramatically induce the global SUMO conjugations (Yoo et al., 2006; Saracoo., et al., 2007). It is necessary to investigate whether the phosphorylation and ubiquitination of SIZ1 are involved in the quick increment of global SUMO conjugation under other environmental changes, especially heat stress, in the future. Recent findings suggest that other components of the SUMOylation pathway, such as SUMO protease ULP1c and ULP1d, are also involved in drought tolerance (Castre et al. 2015). It is worth investigating how these components correlate with the SnRK2-SIZ1 module to control SUMOylation dynamics in the future.

Previous studies have suggested that COP1 directly catalyzes the ubiquitination of SIZ1, leading to the regulation of SIZ1 protein stability (Kim et al., 2016). In the *cop1-4* mutant, the AtSIZ1 protein level and global SUMO conjugation levels were found to be higher under stress conditions when compared to wild-type seedlings (Kim et al., 2016). Light-dependent translocation of COP1 causes SIZ1 proteins to be more stable under light conditions than in darkness (Kim et al., 2016). Under light conditions, COP1 proteins are mainly located in the cytosol, which prevents them from mediating the degradation of nucleus-located SIZ1 (Stacey et al., 1999; Von Arnim and Deng, 1994). In darkness, COP1 translocates from the cytosol into the nucleus to mediate the ubiquitination and degradation of SIZ1. Our data showed that SIZ1 is more stable in light conditions than in darkness in wild-type seedlings (Figure 7B). Surprisingly, the SIZ1 protein level was undetectable in *snrk2-triple* mutant seedlings in the dark, suggesting a primary role of SnRK2s in stabilizing SIZ1 proteins. Consistent with this piece of data, we were unable to detect the SIZ1^S820A^ protein with the abolishing mutation of SnRK2 phosphorylation site, even under light conditions with 12-hour treatment of MG132. Thus, SnRK2-mediated Ser820 phosphorylation is required for the stabilization of SIZ1, even under unstressed dark conditions. Taken together, our findings indicate that SnRK2-mediated phosphorylation stabilizes SIZ1, while a COP1-independent degradation pathway negatively regulates SIZ1 stability. These processes interplay to determine SIZ1 accumulation and promote an increase in global SUMO conjugations in response to osmotic stress and ABA treatment. However, we failed to obtain *snrk2.2/2.3/2.6/siz1-2* or *snrk2.2/2.3/2.6/cop1-4* quadruple mutants, suggesting that this regulation may also be important for embryo establishment and early development. Other environmental challenges, such as high temperatures, also dramatically induce global SUMO conjugation (Yoo et al., 2006; Saracco et al., 2007). It the future, it needs more efforts to clarify the COP1-independent SIZ1 degradation machinery and how phosphorylation of SIZ1 abolishes this degradation machinery upon stresses.

In Arabidopsis, three SnRK2s, SnRK2.2, SnRK2.3, and SnRK2.6 mediate ABA signaling and all the ten SnRK2s are central components in osmotic stress signaling. Though both ABA dependent SnRK2s and ABA-dependent SnRK2s interact with and phosphorylate SIZ1 in the transient assay in tobacco and the *in vitro* kinase assays (Figure 3 and 4), dysfunction of three ABA-activated SnRK2s in *snrk2-triple* is sufficient to abolish Ser820 phosphorylation and result in undetectable SIZ1 protein in dark (Figure 6). During the in vitro kinase assay, SnRK2.4 exhibited stronger phosphorylation of SIZ1 than SnRK2.6, possibly due to SnRK2.4 having stronger interaction with SIZ1, than that of SnRK2.6 in the *in vitro* system. Thus, three ABA-dependent SnRK2.2, SnRK2.3, and SnRK2.6, may have a dominant function in controlling SIZ1 stability *in vivo* in both ABA and osmotic stress conditions. SIZ1 targets transcription factors like MYB30, ABI5, and CaDRHB1, to regulate drought stress and ABA responses (Cheong et al., 2009; Zheng et al., 2012; Joo et al., 2022). The SIZ1 may play a crucial role downstream of SnRK2s in regulating multiple stress adaption processes. To gain a better understanding of these processes, it would be worth identifying more SIZ1 targets and their functions by high-throughput proteomics method. Additionally, another SUMO E3 ligase, HYP2, is also known to participate in stress response (Zhang et al. 2013). The mannitol-induced global SUMO conjugations are also impaired in *hyp2* mutant, and our preliminary data shows that HPY2 also can be phosphorylated by SnRK2s (Supplemental Figure 3D). While this study only focused on the SnRK2-SIZ1 module, it would be interesting to investigate how SnRK2s regulate HYP2 activity and its role in stress response in the future.

In summary, this study reveals the critical role of SnRK2-mediated phosphorylation in regulating global SUMO conjugations by enhancing its stability. We present evidence suggesting a crosstalk between phosphorylation and SUMOylation in osmotic stress and ABA signaling, underscoring the intricate interplay of PTMs in plants. These findings deepen our understanding of the molecular mechanisms underlying plant responses to stress and lay the groundwork for future research on this topic.

## Methods and Materials

### Plant materials

The *Arabidopsis thaliana* Columbia-0 (Col-0) ecotype, *siz1-2* (*SALK_065397*), *hpy2-2* (*SAIL-77-G06*), *snrk2-triple* (*snrk2.2/3/6*), *snrk2-decuple* (*snrk2.1/2/3/4/5/6/7/8/9/10*)(Fujii et al., 2011), *cop1-4,* and *His-H89R-SUM1* (Miller et al., 2010), were used in this study. The seeds were surface-sterilized and grown on half-strength Murashige and Skoog (MS) Agar plates under 16 h light/8 h dark conditions at 23 °C after incubation at 4 °C for 3 days. For full dark growth, the seeds were first exposed to 6h of light to induce germination, and then transferred to continues dark conditions and grown for five days at 23 °C.

To generate transgenic plants expressing *SIZ1-FLAG* under the control of its native promoter, the genomic DNA fragment of SIZ1 with its native promoter sequences were cloned into pEarleyGate302 or pGWB vector, and the resulting constructs were transformed into *siz1-2* using the Agrobacterium-mediated floral dip method. Site-directed mutagenesis was introduced into the pEarleyGate302-SIZ1 or pGWB-SIZ1 constructs using specific primers listed in Supplemental Dataset 2.

### Phenotypic analysis

To perform germination assays, sterilized seeds were sown on 1/2 MS plate containing 0.7% agar and the indicated concentration of ABA, mannitol, NaCl, or PEG, respectively. Radicle emergence was analyzed 72 h after the plates were transferred to 23 °C under a 16 h light/8 h dark cycle photoperiod. Photographs of seedlings were taken at indicated times after transfer to light.

For growth assays, sterilized seeds were grown vertically on 1/2 MS plate with 0.7% agar for 4 days and transferred to medium with or without the indicated concentration of ABA, NaCl, mannitol, or PEG. Root length was measured at the indicated days.

### Generation of anti-SIZ1 antibody

The anti-SIZ1 antibody was generated by ABcloneal using a 6His-SUMO tagged SIZ1 fragment (aa 587–873) expression and purified from E*. coli* strain BL21 (DE3). After removing the HIS-SUMO tag by a SUMO protease ULP1a, the SIZ1 fragment were used as an antigen to immunize rabbit and generate the polyclonal antibody.

### Immunoblot analysis

To determine the levels of SUMO conjugations and SIZ1 concentration, total proteins were extracted from Arabidopsis seedlings using a buffer composed of 50 mM Tris-HCl (pH 8.0), 150 mM NaCl, 1 mM EDTA (pH 8.0), 1 mM DTT, 20 mM NEM, 1% TritonX-100, and 1 × complete protease inhibitor mixture (Roche). A total of 30 µg protein extracts were mixed with 4 × SDS loading buffer (250 mM Tris-HCl pH 6.8, 8% sodium dodecyl sulfate, 0.2% bromophenol blue, 40% glycerol, 20% β-mercaptoethanol) and boiled at 95 °C for 10 min. The proteins were then separated in 8-12% SDS PAGE gels and blotted onto polyvinylidene fluoride (PVDF) membranes. The blots were probed with primary antibodies against SUMO1 (Abcam) at a dilution 1;7500, His (Abmart) at a dilution 1:2500, MBP (NEB) at a dilution 1:5000, and Actin (ABclonal) at a dilution 1:10,000. The primary antibodies were detected with either secondary anti-rabbit or anti-mouse antibodies conjugated with horseradish peroxidase and enhanced chemoluminescence reagent (ShengEr).

### Split luciferase (LUC) complementation assay

Full-length coding sequences of SIZ1 and SnRK2s were cloned into pCAMBIA1300-cLUC and pCAMBIA1300-nLUC vectors, respectively, to perform a Split-LUC complementation assay. The assay was conducted by transient expressing the vectors in tobacco leaves through Agrobacterium-mediated infiltration. After two days of infiltration, luciferase activity was detected with a CCD camera and XenoLight D-Luciferin – K^+^ Salt Bioluminescent Substrate (PerkinElmer).

### GST-pull down assay

For the GST pull-down assay, the full-length coding sequences of SnRK2.4 and SnRK2.6 were cloned into pGEX-4T-1 vectors, while the CDS of SIZ1-C (residues 435-873) fused with an N-terminal-His tag was cloned into pMAL-p2x vectors. The resulting constructs were transformed into E. *coli* BL21 (DE3) and expression was induced for 10-12 h at 20 °C with 0.3 mM isopropylthio-β-galactoside. Equal amounts of GST, GST-SnRK2.4, GST-SnRK2.6 and His-MBP-SIZ1-C were mixed separately with high-affinity GST resins (GE Healthcare) in a pull-down buffer containing 50 mM Tris-HCl (pH 7.4), 120 mM NaCl, 5% glycerol, 0.5% Nonidet P-40, 1 mM phenylmethylsulfonyl fluoride, and 1 mM DTT, and incubated at 4 °C for 2 h. The beads were then washed five times with wash buffer (50 mM Tris-HCl, pH 7.4, 120 mM NaCl, 5% glycerol, and 0.5% Nonidet P-40) and the bound proteins were eluted by boiling the beads with SDS containing sample buffer for 10 min. The eluted proteins were separated by 10% SDS-PAGE and immunoblotted with anti-GST(ABclonal) and anti-MBP (NEB) antibodies.

### Yeast-two-hybrid assay

To investigate protein interactions between SIZ1 and SnRK2s, the full-length CDS of SIZ1 was cloned into pGADT7 vectors and co-transformed with members of pGBKT7-SnRK2.4 and SnRK2.6 into Saccharomyces cerevisiae AH109 cells. Positive colonies were identified on yeast SD medium lacking Leu and Trp or SD medium lacking Leu, Trp, His, and the self-activation was reduced by adding 3-amino-1,2,4-triazole (3-AT).

### Bimolecular fluorescence complementation assays

The constructs SnRK2.4-YNE and SIZ1-YCE or SnRK2.6-YNE and SIZ1-YCE were transiently expressed in N. benthamiana leaves by transfection. After 2-day infiltration, YFP fluorescence signal was detected by confocal microscope.

### In vitro kinase assay

The purified recombinant proteins His-SIZ1-N, His-SIZ1-M, and His-MBP-SIZ1-C, were incubated with GST-SnRK2.4 in 25 µl reaction buffer (25 mM Tris-HCl, pH 7.4, 1 mM DTT, 12 mM MgCl2, 1 μCi [γ-^32^P] ATP and cold ATP) at 30°C for 30 min. The reaction was then stopped by adding SDS loading buffer. The proteins were separated by 10% SDS-PAGE gel and detected by autoradiograph.

### Quantitative RT-PCR Analysis

Total RNA was extracted from seedlings using the RNeasy Plant Mini Kit (QIAGEN) and treated with RNase-free DNase to remove any traces of genomic DNA. One microgram of total RNA was reverse transcribed using the iScript cDNA Synthesis kit (Bio-Rad). Real-time PCR was performed on CFX97 Touch Real-Time PCR Detection System (Bio-Rad) with 1µl cDNA, 1µl forward and reverse primers, and 10 µl iQ SYBR Green Supermix (Bio-Rad) in 20 µl reaction volume. The actin 2 gene was used as a control. The primers used in qRT-PCR are listed in Supplemental Dataset 2.

### LC-MS/MS analysis

The LC-MS/MS analysis was carried out following the protocol described in Lin et al. (2020). Recombinant proteins were produced as previously mentioned, and the phosphorylation reaction was performed using GST-SnRK2.4 as the kinase and GST-SIZ1 as the substrate. The proteins were trypsin digested at a 1:50 trypsin: protein ratio [w/w]) overnight at 37°C, and the resulting phosphopeptides were enriched, and used for LC-MS/MS analysis with an Orbitrap Fusion Tribrid Mass Spectrometer (Thermo Scientific). Proteome Discoverer (Thermo Scientific; version 2.2) was used to analyze the MS/MS spectra, and the phosphorylated peptides were manually inspected to confirm the assignment of phosphorylation sites.

For identification of SUMOylation and Ubiquitination of SIZ1 protein, the 10-day-old *His-H89R-SUM1* seedlings (Miller et al., 2010) were first treatment with 20 μM MG132 for 6 h, followed by treatment with 50 μM ABA for 1 h. 5 g frozen tissue were ground to fine powder and total proteins were extracted with extraction buffer containing 50 mM Tris-Cl (pH 8.0), 150 mM NaCl, 2 mM EDTA, 20 mM NEM, 0.1% TritonX-100, 0.2% Nonidet P-40, 0.6 mM PMSF and 1× complete protease inhibitor mixture (Roche). After removing of cell debris by centrifug at 16,000g at 4°C for 20 min, the protein extract was incubated with 30 μL Protein A Agarose beads (Millipore), 10 μL anti-SIZ1 antibody with rotation at 4°C for 4 h.

The beads were then washed with extraction buffer once, followed by wash buffer (50 mM Tris-HCl, pH 8.0, 150 mM NaCl, 2 mM EDTA, pH 8.0, 20 mM NEM) for five times, and the on-beads trypsin digestion was performed. The resulting peptides were analyzed using an Orbitrap Fusion Tribrid Mass Spectrometer (Thermo Scientific), and the MS/MS spectra were compared against the protein sequences of SIZ1 using Thermo Proteome Discoverer (version 2.2).

### Parallel Reaction Monitoring

For Parallel reaction monitoring, proteins were extracted from seedlings according to the protocol described by Wang et al. (2020). Briefly, wild type and *snrk2* mutant seedlings with or without ABA and mannitol treatment were ground to a fine powder in liquid nitrogen and lysed in 6 M guanidine hydrochloride comprising of 100 mM Tris-Cl (pH 8.5), ethylenediaminetetraacetic acid free protease inhibitor mixture (Roche) and phosphatase inhibitor mixture (Sigma-Aldrich). The protein extracts were then reduced and alkylated with 10 mM Tris-(2-carboxyethyl) phosphine proteins and 40 mM chloroacetamide. Subsequently, the proteins were precipitated using methanol-chloroform, and the pellets were dissolved in 12 mM sodium deoxycholate (SDC) and 12 mM sodium lauroyl sarcosinate (SLS). The concentration of protein was determined by Bicinchoninic Acid assay Kit (Thermo Fisher Scientific). Diluted and digested with Lys-C (Wako) at 37°C for 3 h followed by trypsin at 37°C overnight.

The phosphopeptides were enriched by IMAC, and then dissolved in 6 μL 2%ACN, 0.2% formic acid and injected 4 μL into an Easy-nLC 1200 (Thermo Fisher Scientific) HPLC system. The eluent was introduced into the mass spectrometer, using a 15 cm in-house packed column (360 µm OD × 75 µm ID) containing C18 resin (2.2 µm, 100 Å, Michrom Bioresources). The mobile phase buffer consists of 0.1% formic acid (buffer A) in water with an eluting buffer of 0.1% formic acid in 80% ACN (buffer B) and the LC flow rate was 300 nL/min. The gradient was programmed to run at 5–25% buffer B for 30 min, 25–45% for 10 min, and 45–90% for 3 min. The sample was then acquired on Thermo Orbitrap Fusion (Thermo Fisher), analyzed under PRM with an isolation width of ± 0.7 Th.

In all experiments, a full mass spectrum was acquired at 30,000 resolution relative to m/z 200 (AGC target 2E5, 300 ms maximum injection time, m/z 150–1500), followed by up to 20 PRM scans at 12,000 resolution (AGC target 1E5, 50 ms maximum injection time), triggered by an unscheduled inclusion list. Higher-energy collisional dissociation was used with 32 eV normalized collision energy. Finally, PRM data were manually curated within Skyline (version 3.5.0.9319).

## Cell-free protein degradation assay

For the cell-free protein degradation assay, twelve-day-old wild type seedlings were harvested and ground to a fine powder in liquid nitrogen. Total proteins were subsequently extracted in degradation buffer comprising 25 mM Tris-Cl, pH 7.5, 10 mM NaCl, 10 mM MgCl2, 5 mM DTT and 10 mM ATP. Cell debris was removed by centrifugation at 12,000 × g at 4°C, and the supernatant was collected. Protein concentration was determined by Bicinchoninic Acid assay Kit (Thermo Fisher Scientific). The His-MBP-SIZ1-C-His and His-MBP-SIZ1-C^S820A^-His recombinant proteins purified from E*. coli* were incubated with an equal amount total protein extracts were at 25 °C for the indicated time points.

## Data availability

The mass spectrometry proteomics data were deposited to the ProteomXchange (PXD040237) and JPST002048 for JPOST (URL: https://repository.jpostdb.org/preview/33548052463f335c633853. Access key: 4522).

## Acknowledgements

This work was supported by the National Key Research and Development Program of China, Grant 2021YFA1300402. We are grateful to Prof. Richard D. Vierstra of Washington University at St. Louis, USA, for kindly providing the *His-H89R-SUM1* seeds, to Prof. Xingwang Deng of Peking University, China, for kindly providing of *cop1-4* seeds.

## Supplemental Materials

Supplemental Figure 1. SIZ1 is involved in osmotic stress.

Supplemental Figure 2. SnRK2s interact with SIZ1.

Supplemental Figure 3. SnRK2s phosphorylate SIZ1.

Supplemental Figure 4. Complementation of mutated SIZ1 in *siz1-2* to rescue wild type.

Supplemental Figure 5. SIZ1 is phosphorylated by SnRK2s at Ser820.

Supplemental Figure 6. MS/MS spectrum of the modification sites of SIZ1.

Supplemental Dataset 1. The SIZ1 SUMOylation and ubiquitination sites in His-H89R-SUM1 seedlings identified by LC-MS/MS.

Supplemental Dataset 2. Primers used in this study.

---

**Supplemental Figure 1.**
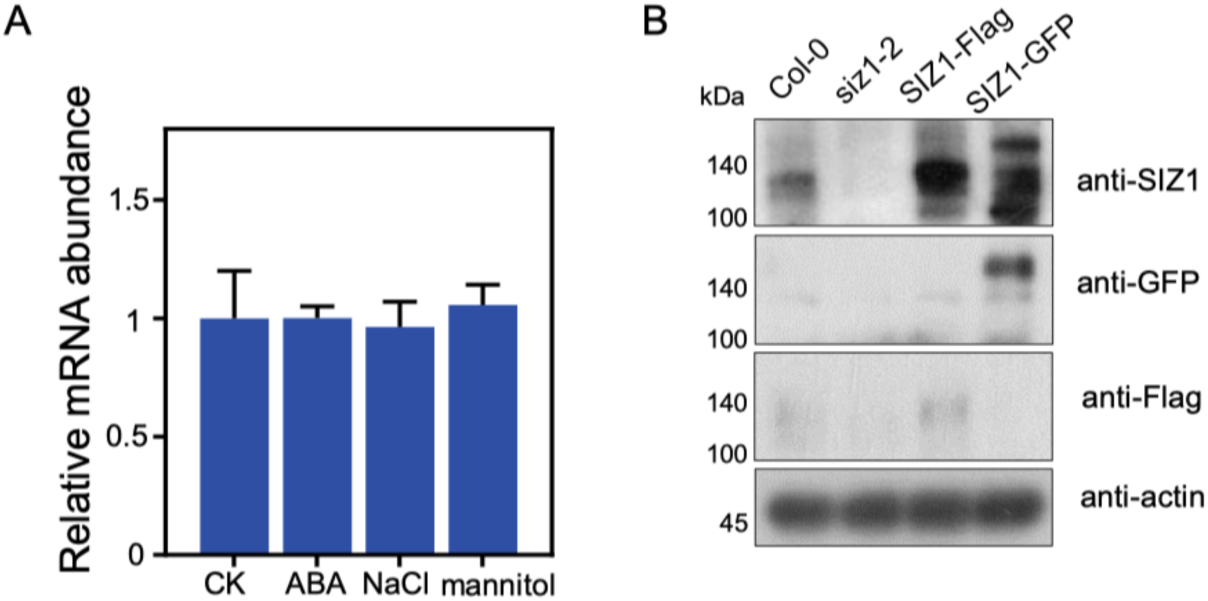
SIZ1 is involved in osmotic stress. **A.** The expression of SIZ1 after indicated time of ABA, NaCl, or mannitol treatments. **B.** The validation of anti-SIZ1 antibody. Immunoblot with anti-SIZ1 antibody (upper) showed the amount of SIZ1 protein in wild type, *siz1-2*, *SIZ1-FLAG* and *SIZ1-GFP* seedlings. Immunoblot with anti-FLAG, anti-GFP and anti-actin was used to show the SIZ1 abundance of transgenic plants and the loading of total proteins in each lane.

**Supplemental Figure 2.**
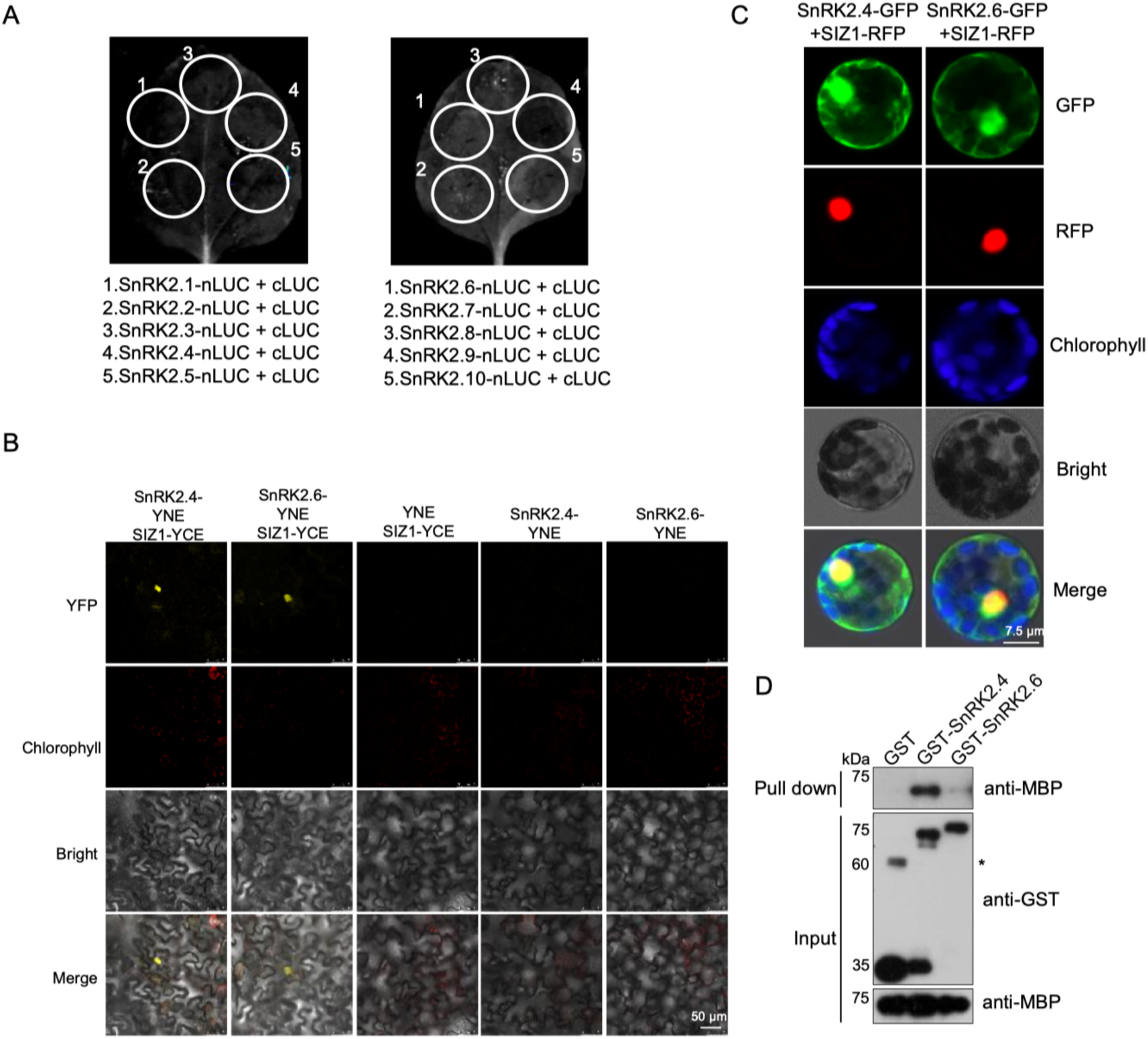
SnRK2s interact with SIZ1. **A.** The cLUC and ten SnRK2s-nLUC were used as negative controls in split LUC assay. **B.** Interaction of SnRK2.4/2.6 and SIZ1 detected by bimolecular fluorescence complementation (BiFC) assays. The signal was detected by confocal microscopy. **C.** Fluorescence of SIZ1-RFP and SnRK2.4-GFP/SnRK2.6-GFP in protoplasts co-transformed assay. D. GST pull-down assay to detect the interaction between SIZ1-C and SnRK2s. The GST-SnRK2.4 and GST-SnRK2.6 were used to pull-down MBP-SIZ1-C. The immunoblot with anti-GST (middle panel) and anti-MBP (bottom panel) antibody were used to show the loading of MBP-SIZ1-C and GST fused SnRK2s. * represents the nonspecific band.

**Supplemental Figure 3.**
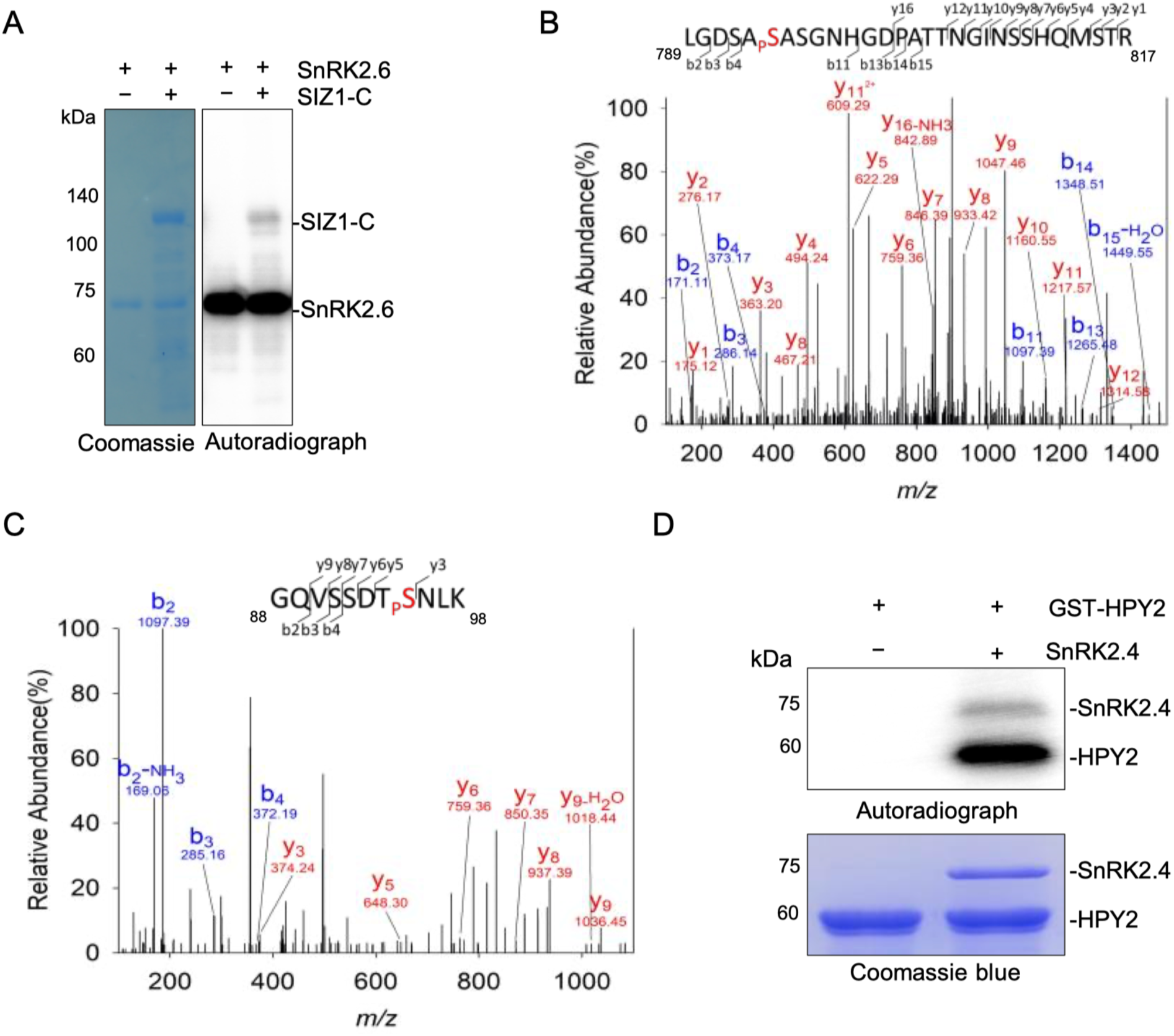
SnRK2s phosphorylate SIZ1. **A**. Recombinant SnRK2.6 was used to phosphorylate recombinant SIZ1-C in the presence of [ψ-^32^p] ATP. Autoradiograph (right) and Coomassie staining (left) show phosphorylation, loading of purified SIZ1-C and SnRK2.6.**B.** The MS/MS spectrum showing that the ^18^O-phosphopeptide GQVSSDTSNLK contains the phosphoserines Ser95 in SIZ1. **C.** The MS/MS spectrum showing that the ^18^O-phosphopeptide LGDSASASGNHGDPATTNGINSSHQMSTR contains the phosphoserines Ser794 and Ser796 in SIZ1. **D**. Recombinant SnRK2.4 was used to phosphorylate recombinant GST-HPY2 in the presence of [ψ-^32^p] ATP. Autoradiograph (right) and Coomassie staining (left) show phosphorylation, loading of purified GST-HPY2 and SnRK2.4.

**Supplemental Figure 4.**
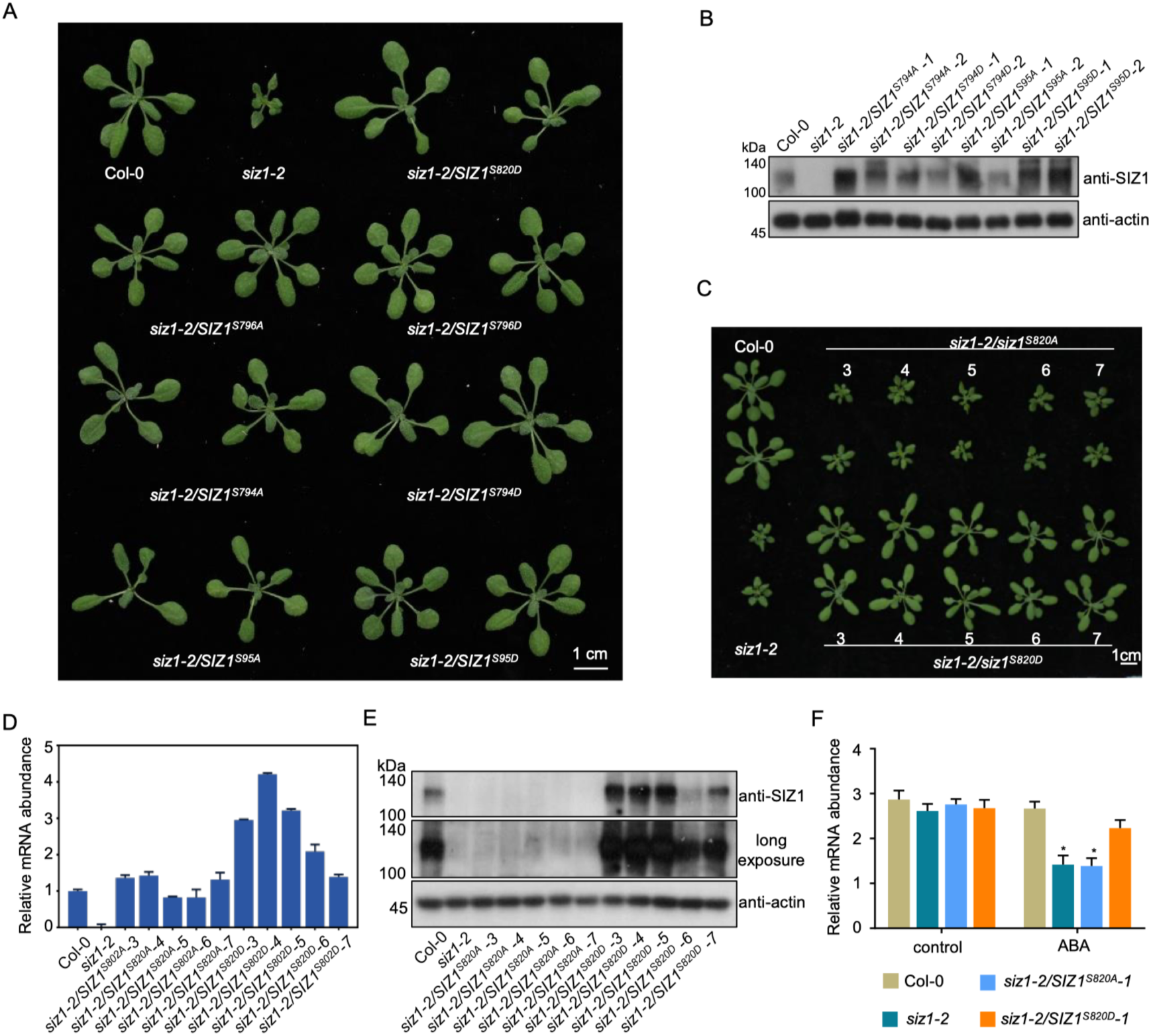
Complementation of mutated SIZ1 in *siz1-2* to rescue wild type. **A.** Photographs of 4-week-old seedlings from the wild type, *siz1-2*, and different transgenic plants. **B.** Immunoblot shows the SIZ1 abundance in wild type, *siz1-2*, and transgenic plants. Immunoblot with anti-actin antibody (bottom) used to show loading of total proteins in each lane. **C**. Photographs of 4-week-old seedlings from the wild type, siz1-2, and *SIZ1^S820A^/siz1-2* and *SIZ1^S820D^/siz1-2* transgenic plants. **D.** Transcription of *SIZ1* genes in wild type, *siz1-2*, *SIZ1^S820A^/siz1-2* and *SIZ1^S820D^/siz1-2* transgenic plants. **E.** Immunoblot shows the SIZ1 abundance in wild type, *siz1-2*, *SIZ1^S820A^/siz1-2* and *SIZ1^S820D^/siz1-2* transgenic plants. Immunoblot with anti-actin antibody (bottom) used to show loading of total proteins in each lane. **F**. Quantitative measurement of the root length of the seedlings after ABA treatment. Values are mean ± SD (n = 15; *, p < 0.05. Student’s t-test).

**Supplemental Figure 5.**
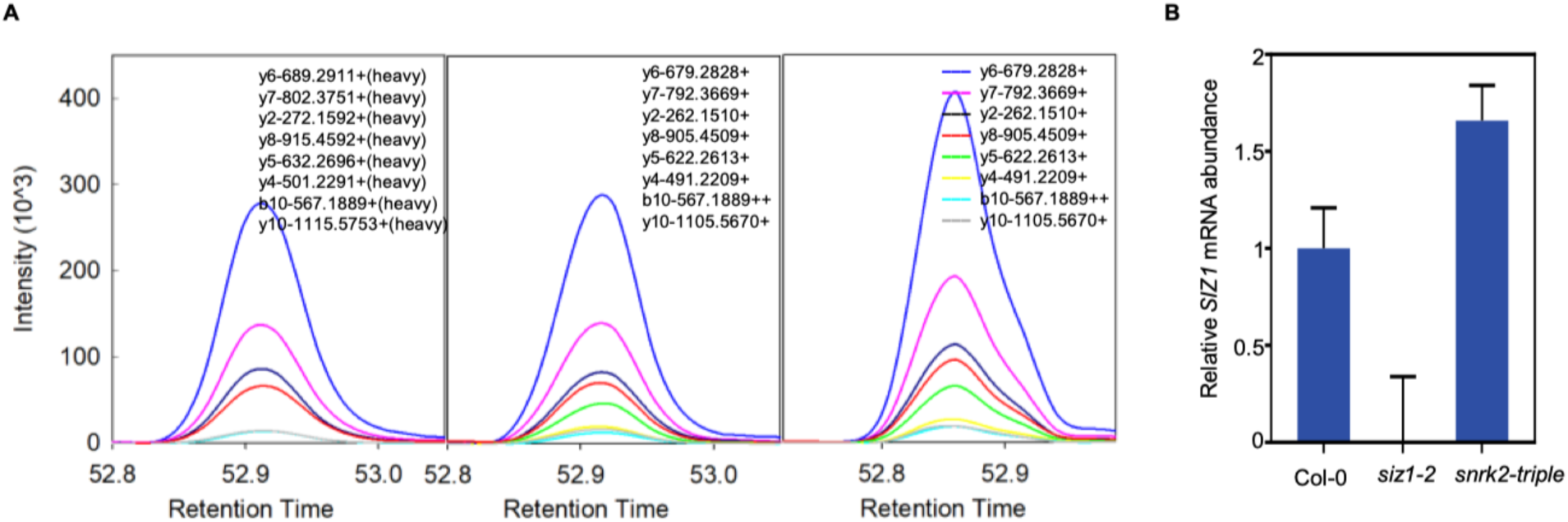
SIZ1 is phosphorylated by SnRK2s at Ser820. **A.** Parallel reaction monitoring (PRM) analysis of the phospho-peptide intensity in wild type with (middle) or without (right) ABA treatment. Graphs displaying chromatograms of fragment ions from the peptide EGpSMDTTTETASLLLGMNDSR (m/z 770.9954, 3+). The heavy isotope labeled-peptide EGpSMDTTTETASLLLGMNDS[^14^C^15^N]R (m/z 774.3315, 3+) was used as internal standards for peptide quantification (left). **B.** Transcription level of *SIZ1* genes in wild type, *siz1-2* and *snrk2-triple* (*, p < 0.05. Student’s t-test).

**Supplemental Figure 6.**
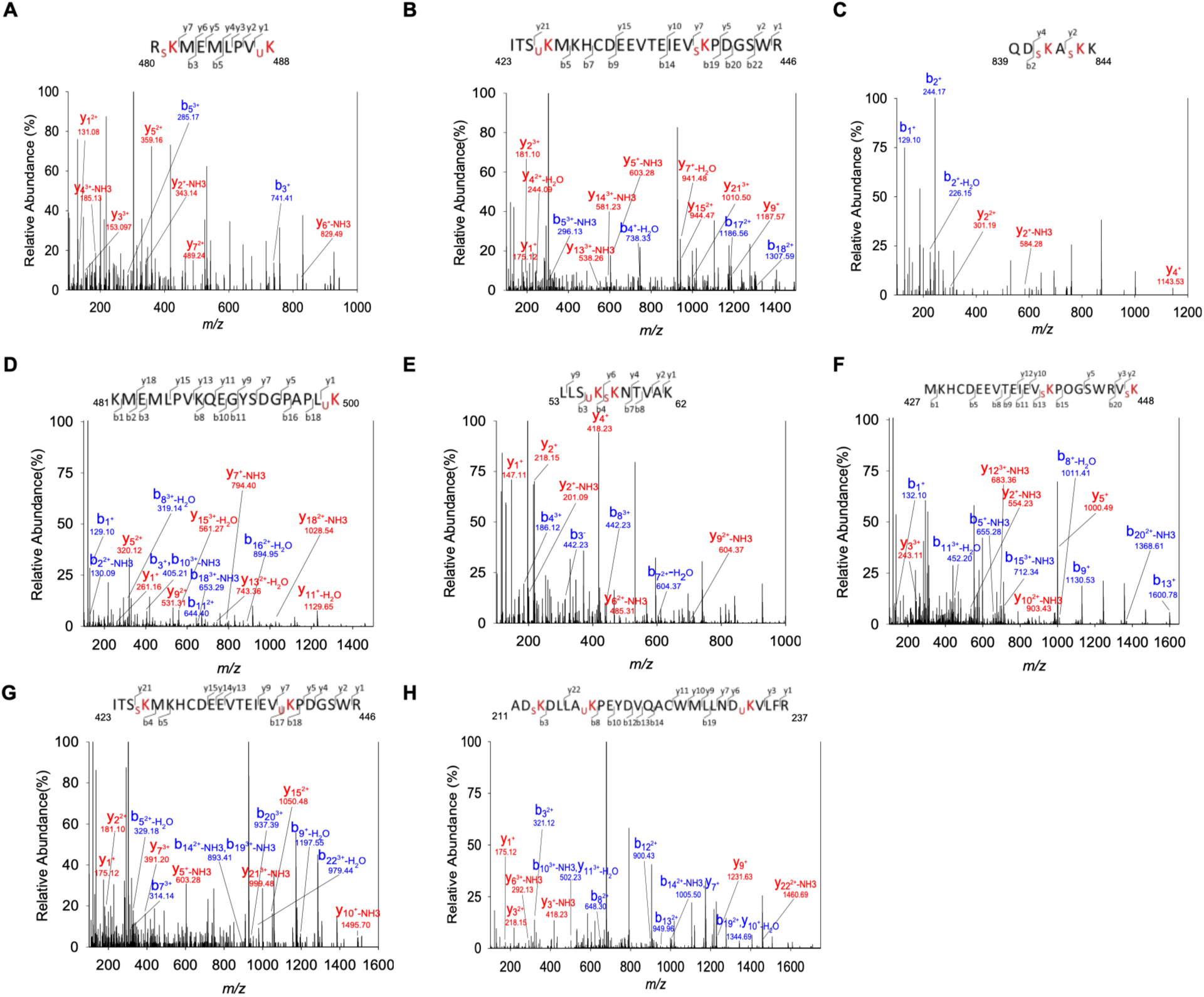
MS/MS spectrum of the modification sites of SIZ1. **A**. MS/MS fragmentation spectra for the K481-SUMO, K488-GG containing peptide. **B**. MS/MS fragmentation spectra for the K426-SUMO, K440-GG containing peptide. **C**. MS/MS fragmentation spectra for the K841-SUMO, K843-SUMO containing peptide. **D**. MS/MS fragmentation spectra for the K500-GG containing peptide. **E**. MS/MS fragmentation spectra for the K56-GG, K57-SUMO containing peptide. **F**. MS/MS fragmentation spectra for the K440-SUMO, K448-SUMO containing peptide. **G**. MS/MS fragmentation spectra for the K440-SUMO, K426-GG containing peptide. **H**. MS/MS fragmentation spectra for the K481-SUMO, K488-GG containing peptide.

